# Discrete Time Modeling of Stock-Recruit Relationships with Life-History Stanzas

**DOI:** 10.1101/2024.03.21.586068

**Authors:** Anna-Simone Frank, Ute Schaarschmidt, Richard D. M. Nash, Sam Subbey

## Abstract

The stock-recruit relationship is a foundational concept in fisheries science, bridging the connection between parental populations (stock) and progeny (recruits). Traditional approaches describe this relationship using closed-form analytical functions, which represent only a restricted subset of the broader class of possibilities. This paper advocates for a novel approach that integrates discrete time modeling with a life-history cycle framework, incorporating distinct stanzas and developmental processes. By breaking down the life cycle into identifiable stages, we capture the step-wise progression of life history traits and the factors influencing recruitment outcomes. Through numerical simulations, we explore the advantages of this approach, including complexity handling, dynamic behavior modeling, and scenario exploration. Our simulation results show that we are able to generate a broad spectrum of stock-recruit relationships (including the traditional ones), which best reflect variability observed in nature. We demonstrate how this framework allows for the identification of critical stages, and integration of various factors that influence recruitment. This holistic approach enhances our comprehension of the intricate interactions shaping stock-recruit relationships and advances our understanding of sustainable population dynamics.

**Highlights:**

- A novel multi-stage life-cycle model is presented.
- Model simulations reveal three different Stock-Recruitment (SR) patterns.
- Our approach contributes to enhanced understanding of SR relationships.

## 1. Introduction

In fisheries science and ecology, stock-recruit (SR) relationships link parental populations to progeny [1]. The SR relation considers the early life history of fishes into a single step and was implemented for stock assessment purposes (see [2]). Understanding this relationship is vital as it provides essential insights into the dynamics of fish populations, allowing us to understand how the abundance of adult spawners relates to the success of recruitment of juvenile fish. Recruitment describes the transition of younger fish (e.g., juveniles) to another developmental stage (i.e., adults) [3]. This understanding is fundamental for making informed decisions on sustainable fishing practices, conservation efforts, and ecosystem management, ensuring the long-term health and resilience of fish populations and the ecosystems they inhabit.

Traditionally, SR-relationships have been described by models that are formulated as closed-form analytical functions [4, 5]. These functions are typically nonlinear and can exhibit a saturating relationship, where the number of recruits increases with increasing progeny abundance but eventually reaches a maximum level or starts to decline as progeny abundance continues to increase. While these functions offer a useful framework, e.g., in the context of management advice [1, 6], there are compelling reasons to argue that they define a restricted subset of possible SR relationship, considering the intricacies and dynamics of natural ecosystems [6].

Some of these reasons are the following. Threshold effects and Allee dynamics [7, 8] are often absent from traditional models. Fish populations, in reality, can exhibit abrupt transitions and non-monotonic responses to changes in stock size, which traditional models may not effectively capture. Similarly, the interplay between density-dependent and density-independent factors [9, 10, 11], introduces complexity that traditional models, which typically emphasize density-dependent processes, may not fully account for. Complex feedback mechanisms, as observed in ecological systems [12, 13, 14], challenge traditional models’ simplicity, leading to non-linear responses in recruitment. Traditional models also overlook the potential for evolutionary responses [15], where fish populations may respond to selection pressures, potentially leading to non-linear stock-recruitment relationships. Additionally, they assume a consistent relationship between stock and recruitment, but fish populations exhibit spatial and temporal variations in recruitment due to environmental conditions and life history strategies [16, 17], which can result in non-monotonic relationships. Human impacts, including fishing pressure and management interventions, disrupt natural stock-recruit relationships [18], introducing abrupt changes or non-linear responses in recruitment. In addition, non-linear growth and reproduction patterns with age and size [19], can lead to dramatic changes in reproductive output, contributing to non-monotonic recruitment responses.

Furthermore, in reality, the stock-recruit relationship encapsulates the early part of the life cycle, which consists of several consecutive stanzas (i.e., development stages) and transitions between them [20]. Therefore, age- and stage-structured models in fisheries are traditionally represented using difference equations, i.e., assuming discrete transitions between life stanzas (see e.g., Fig. 1), uniform time steps across all life stanzas, and a positive time delay between spawning and recruitment [21]. These assumptions are consistent with the fisheries literature [22, Chapter 5]. In addition, there are various ways to define recruitment, and it may be associated with juveniles or maturity. This knowledge has motivated the development of several multistage models, e.g., [23, 24], as well as life cycle approaches that encompass two life stages, namely, pre-recruits and adults [25, 26].

**Figure 1:**
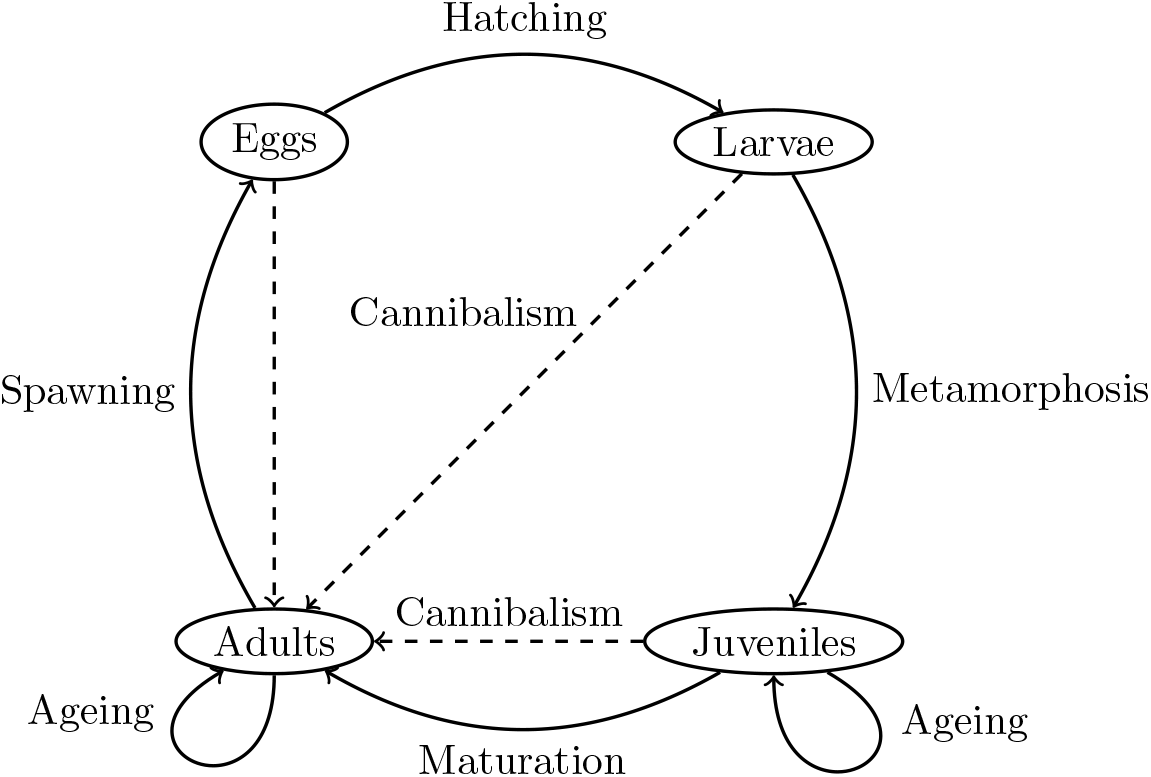
Illustration of the life-history cycle of a fish with four stages. Arrows represent the principal developmental processes (which cause the transition from one life-history stage to the next) and cannibalism as an example of feedback mechanisms (From [39]). Loops represent transitions between two age classes within one stage.

The existing models have mainly focused on the well known SR-functional relationships, such as Ricker-, Beverton-Holt or Hokey stick functions (see e.g., [24]). However there is a full spectrum of SR-relationships that are observed in nature, but that have not been confirmed computationally. For example, non-linear, periodic, oscillatory and even chaotic relationships have been observed [27, 28, 29, 30, 31, 32, 33, 34]. This underscores the need for more comprehensive and flexible modeling approaches that considers each stage and state transition in the fish species’ life cycle, to gain a comprehensive understanding of the SR-relationship and its emergent patterns. Whether each observed SR-relationship can be represented by a closed form analytical function, as in the case of Ricker- and Beveron-Holt functions, has to be investigated.

To our knowledge, few attempts have been reported in the literature to model the entire life cycle comprehensively [35, 36, 37, 38]. This is due to the complexity involved and uncertainty regarding the actual relationship and influential factors. However none of these articles used a life cycle model to infer the SR-relationship.

A recent attempt has been made in [39]. The authors present a general discrete time multi-stage model (DTMM) approach of the complete fish lifecycle. They investigate the theoretical necessary and sufficient conditions for the existence of a functional SR-relationship in its generic form. The approach is based on the assumption that spawning and transition to the juvenile stage occur within one time step and juveniles and adults age by one in every simulation time step (if surviving). This is justified by typical stage durations of several fish populations described in [40].

This paper implements the DTMM framework by [39] to model the entire life cycle of fish. The primary objective is to explore possible emergent SR-relationships (“pattern”) of the DTMM approach through numerical simulations. The secondary objective is to investigate whether the identified SR-relationships have a closed-form functional representation.

The article is organized in the following way. In Section 2, we present the model assumptions and the discrete-time multi-stage model (DTMM) that will be implemented. In Section 3, we present aspects of parameter estimation, model implementation and validation as well as the numerical scenarios. Section 4 presents the presents the main results, and finally we discuss the results and draw a conclusion in Section 5.

## 2. Life-cycle modeling approach

### 2.1 Biological assumptions and examples

As in [39], our life-cycle model describes the life-history cycle illustrated in Figure 1 and we adopt the following modeling assumptions.

We assume that the life-history cycle includes four stages: eggs, larvae, juveniles, and adults. Juvenile stages and adult stages may consist of several age classes. A proportion of adults, spawners, contribute to egg production. Surviving eggs transition into the larval stage, and may survive to become juveniles. Survival rates also determine the transitions of juveniles as they age. At a given time step/age, surviving juveniles are recruited into the adult stage. This defines recruitment for our model. The adult stage consists of several age classes, and all of them could include spawners. Concerning the relationships between stages, we assume:

**B1:** Egg production depends on fecundity and the total number of spawners (as described e.g., in [41, Chapter 7]

**B2:** Survival rates may be affected by processes such as cannibalism, food availability, and competition (as described e.g., in [41, Chapter 7])

More specifically, we assume that (i) juveniles within each age group compete for the same food resource and habitat and (ii) the adult population could cannibalize eggs, larvae and juveniles.

These assumptions are summarized in the next subsection in form of model equations.

### 2.2 The life cycle model (DTMM)

The age- and stage-structured population dynamic model defined by [39] describes births, survivals, and transitions from one stanza to the next. The stages are eggs (*e*), larvae (*l*), juveniles (*j*), and adults (*a*). Age classes 0, …, *m* −1 represent juveniles and age classes *m*, …, *n* adults. The symbols in brackets and 0, …, *n* are used as indices. The transitions from one stage or age class to the next are illustrated in Figure 1. The model represents the age- and stage-structured population using difference equations. Herein, it is assumed that spawning and transition to the juvenile stage occur within one time step and juveniles and adults age by one in every simulation time step (if surviving). The general discrete time multi-stage model is included in appendix Appendix A. There, we summarize the model assumptions and equations, as well as conditions and theorems from [39] that are relevant for Sections 3–5.

Based on the general model equations in (A1–A7) and to be able to simulate the population numerically, we make the following additional assumptions:

**B3:** Survival of eggs is an exponential function of the numbers of adults.

**B4:** Survival of larvae is proportional to the probability of surviving competition and the probability of surviving cannibalism. Competition is proportional to the number of individuals and survival represented by a logistic curve (as described e.g., by [42, Chapter 2.1,2.4]. Following [4], we assume the instantaneous mortality rate due to cannibalism to be proportional to the number of predators. Survival of cannibalism is an exponential function of the numbers of adults.

**B5:** As for larvae, survival of juveniles is proportional to the probability of surviving competition (represented by a logistic curve) and the probability of surviving cannibalism (represented by an exponential function).

**B6:** Survival of adults is the product of the probability of surviving fishing and natural mortality. The two mortality rates of adults are functions of age. We assume the natural mortality rate to be a bath-tub function of age (i.e., with increasing age, mortality first decreases, then stays approximately constant, and finally increases). This pattern represents three phases: initial, stable, and death by senescence [43, 44]. We assume fishing mortality increases with age. Logistic functions of age may represent both availability and vulnerability (to fishing gear) [45].

**B7:** Fecundity is a potentially asymmetric and dome-shaped function of age, i.e., it has one local maximum in the interior of its domain. This assumption arises from the consideration that the joint effects of spawning frequency and spawning duration are represented by a dome-shaped function [46].

To define the DTMM, we use the notation from Table 1. Based on the general discrete time multi-stage model by [39] under additional assumptions **B1–B7**, our DTMM is given by (1)–(10).

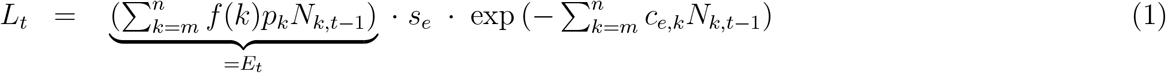

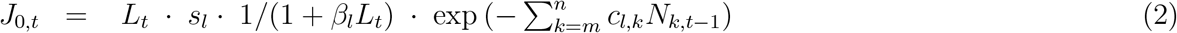

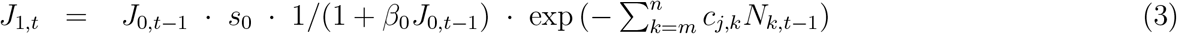

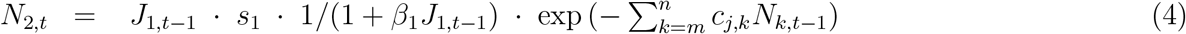

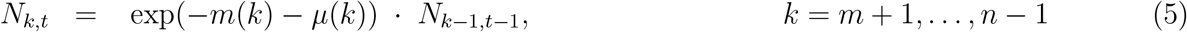

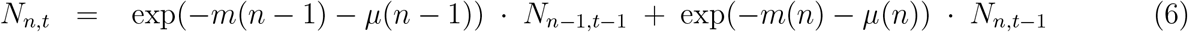

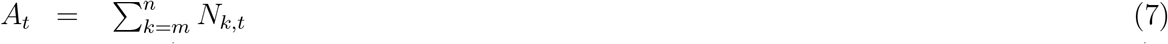

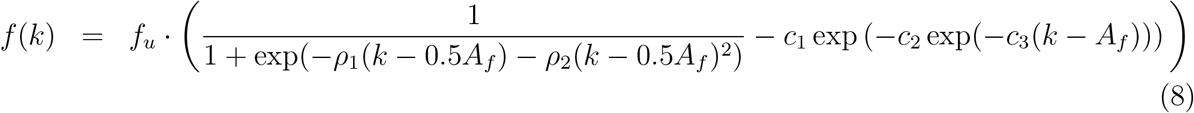

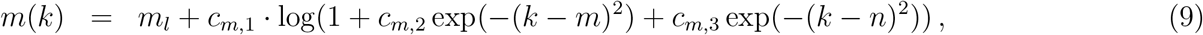

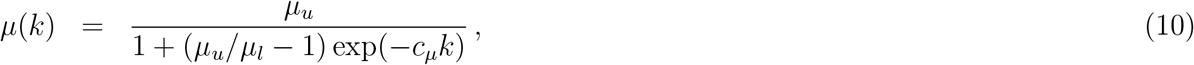

**Table 1:** Nomenclature for the discrete time multi stage model.

| **Variables and indices** |  |
| --- | --- |
| $\mathcal{S} = \{e, l, j, a\}$ | Set of life stages: eggs ($e$), larvae ($l$), juveniles ($j$), adults ($a$) |
| $m \in \mathbb{N}_0$ | Number of juvenile age classes |
| $n \in \mathbb{N}_0$ | Total number of age classes |
| $k \in \{0, \dots, m, \dots, n\}$ | Indices of age classes |
| $t \in \mathbb{N}_0$ | Simulation time |
| $E_t \in \mathbb{R}_{\geq 0}$ | Number of eggs at time $t$ |
| $L_t \in \mathbb{R}_{\geq 0}$ | Number of larvae at time $t$ |
| $J_{k,t} \in \mathbb{R}_{\geq 0}$ | Number of juveniles in class $k = 0, \dots, m - 1$ at time $t$ |
| $N_{k,t} \in \mathbb{R}_{\geq 0}$ | Number of adults in class $k = m, \dots, n$ at time $t$ |
| $R_t = N_{m,t}$ | Number of individuals included in stage $m$ at time $t$ |
| $A_t$ | Number of adults at time $t$ |
| **Probabilities and functions** |  |
| $f : \mathbb{N} \rightarrow \mathbb{R}$ | Annual fecundity of adult fish as a function of age |
| $s_i \in [0, 1]$ | Probability of surviving stage $i \in \{e, l, 0, \dots, m - 1\}$ in a time step in case there is no competition and no cannibalism |
| $\mu : \mathbb{R}_{\geq 0} \rightarrow (0, 1]$ | Fishing mortality rate of adults |
| $m : \mathbb{R}_{\geq 0} \rightarrow (0, 1]$ | Natural mortality rate of adults |
| **Parameters** |  |
| $p_k \in (0, 1)$ | Proportion of adult female population in class $k = m, \dots, n$ |
| $c_{i,k} \in \mathbb{R}_{\geq 0}$ | Coefficient for cannibalism attributed to fish of stage/age $i \in \{e, l, j\}$ per adult of age $k = m, \dots, n$ |
| $\beta_i \in \mathbb{R}_{\geq 0}$ | Coefficient for competition attributed to fish of stage/age $i \in \{l, 0, \dots, m - 1\}$ per individual in the same stage/age |
| $f_u \in \mathbb{R}_{\geq 0}$ | Scaling constant for fecundity |
| $\rho_1, \rho_2 \in \mathbb{R}$ | Parameters for fecundity |
| $A_f, c_1, c_2, c_3 \in \mathbb{R}_{\geq 0}$ | Non-negative parameters for fecundity |
| $m_l \in \mathbb{R}_{\geq 0}$ | Lower bound for natural mortality of adults |
| $c_{m,1}, c_{m,2}, c_{m,3} \in \mathbb{R}_{\geq 0}$ | Parameters for natural mortality of adults |
| $\mu_l, \mu_u \in \mathbb{R}_{\geq 0}$ | Lower and upper bound for fishing mortality of adults |
| $c_\mu \in \mathbb{R}_{> 0}$ | Parameter for fishing mortality of adults |

Total egg production at time *t* is a function of the numbers of individuals in the adult population and is defined as the sum of eggs produced over all age classes. Within the same time step, the number of eggs that survive becomes the number of larvae. Following **B1** and **B3**, numbers of eggs and larvae are thus given by (1). Still, in the same time step as egg production, the initial juvenile population of age 0 transitions from the larvae stage. The number of juveniles of age 0 is defined in (2) and depends, as described in **B4**, on the number of larvae and the adult fish population. We assume that there exist two age classes of juveniles (i.e., *m* = 2). Following **B5**, juveniles of age 1 have survived cannibalism and competition with other juveniles of age 0 and their number is given by (3). Survival of juveniles of age 1 also follows assumptions **B5** and recruitment to adults of age 2 is given by (4). Survival rates of adults (5) are linked to fishing and natural mortality as described in **B6**. The oldest age class of adults presented in (6) may consist of several age classes *k* ≥ *n*. The total number of adults (7) is in the following also referred to as spawning stock.

Our definition of fecundity (8) is a generalization of fecundity as a function of age as presented by [46]. Following [46], we aim for a dome-shaped function for fecundity. However, as we want the graph of *f* to be asymmetrical with positive asymptotic value lim_*k*→∞_ *f* (*k*), we add an exponential term. Our definitions of natural and fishing mortality of adults (9)–(10) are based on assumption **B6**. The logistic equation (10) has lower bound *μ*_*l*_ ≥ 0 and upper bound *μ*_*u*_ ≥ 0.

## 3. Numerical simulation approach

In this section, we present the numerical approach that is used to generate a broader range of SR-relationship patterns that are known to exist.

We base our primary analysis on numerical simulations, as they excel in handling the complexity of intricate relationships, capturing feedback loops and complex interactions more realistically than simpler analytical models [47]. They also enable the modeling of dynamic behaviors and emergent properties, accounting for, among other things, interactions between different life-history stages, environmental fluctuations, and variable population sizes [47].

To simulate SR-relationships with the DTMM described in Section 2, we first estimate model parameters from available data. Then, we will describe the Paulik diagram as tool for model validation, i.e., pattern recognition, and finally we will present the different numerical scenarios.

### 3.1 Parameter estimation from available data

Estimating parameters in population dynamic models is challenging due to the following reasons (adapted from [21, p. 8, 68]): Models may incorporate several species or development stages as well as environmental factors with temporal and spatial variations at several scales. Such diverse information in time, space and scale is often however not represented in available data that, for example, include measurement of numbers of recruits and adults at several time points. This leads to the fact that estimated model parameters from data are often highly uncertain. Another result of this situation is that implausible model results have been reported especially when parameters in population dynamic models were estimated from data. As a consequence, the fisheries literature has adopted a heuristic approach, where for example, some parameters are fixed (based on e.g., literature reports), while others are successively changed.

Acknowledging this challenge of parameter selection, we present example parameter values that serve as illustrative cases. Following common practice, we use an heuristic approach, where some parameters are taken directly from the literature, whiles others were estimated using optimization procedures.

As basis for these parameter estimates, we used data of North Sea cod, since this species is well documented. For this stock, data are available from 1963 and for age classes 1 to 7+, with the plus age class including ages greater or equal to 7 [48]. In particular, the numbers of fish of ages 1 to 6 at time 0 are numbers of North Sea cod for the year 1963 [48, Table 14.9]. We include more than 7 age classes and our guesses for initial values for *N*_7,0_, …, *N*_20,0_ are derived from the number of individuals in age class 7+ in year 1963. We consider two age classes of juveniles, i.e., *m* = 2, and assume that *n* = 20.

The natural mortality terms of adults are very uncertain and are therefore estimated from available data. That means that the values for *m*_*l*_, *c*_*m*,1_, *c*_*m*,2_, and *c*_*m*,2_, which describe natural mortality of adults, are obtained from minimization of the least squares error between *m*(*k*) given by (9) and natural mortality data for North Sea cod [48, Table 14.5.b]. Here, we employ the Levenberg-Marquardt method for parameter estimation as implemented in [49]. For the remaining parameters, we use synthetic values. Model parameters and initial conditions used in the numerical experiments are presented in Subsection 3.3.

### 3.2 Validation of stock-recruitment relationships

To demonstrate possible existing functional structures i.e., patterns, of the resulting SR-relationship and to study their connection to the underlying assumptions, we use Paulik diagrams [50, 51]. A Paulik diagram visualizes the relationship between four separate life stages in one figure. The underlying idea is to illustrate the graph of three functions *g*_1_, *g*_2_ and *g*_3_ in quadrants IV, III, and II of a Paulik diagram. The composition *g*_3_ ∘ *g*_2_ ∘ *g*_1_ is presented in the first quadrant, and represents their combined dynamic.

To create a Paulik diagram [50], we need to use four of the six life stages that are included in our DTMM (see Section 2). We choose to focus on the stages of recruitment, adults, larvae, and juveniles of age 0. The reason for not including eggs is that the relationship between adults and eggs given in (1) is linear, while other relationships are in general non-linear. The inclusion of *J*_0,*t*_, instead of *J*_1,*t*_, is based on the assumption that competition and cannibalism might be more relevant for younger fish than for older ones (e.g.,[52, 53, 54, 55]). Thus, *J*_0,*t*_ is considered a more critical stage than *J*_1,*t*_ and therefore included in the Paulik diagram.

For the life-cycle considered in this article, let *A*_*t*_ (adults), *L*_*t*_ (larvae), *J*_0,*t*_ (juveniles of age 0) and *R*_*t*+2_ (recruits) (see Table 1) for *t* = 10, …, 50 be given by the DTMM (1)–(10). The quadrants of the Paulik diagrams in the result section represent the following relationships:

- Quadrant IV: *A*_*t*_ on the x-axis and *L*_*t*_ given by (1) on the y-axis, i.e. the total number of adults and the number of larvae spawned by them and surviving the egg stage.
- Quadrant III: *L*_*t*_ on the y-axis and *J*_0,*t*_ described by (2) on the x-axis.
- Quadrant II: *J*_0,*t*_ on the x-axis and the number of recruits *R*_*t*+2_ = *N*_2,*t*+2_ spawned in the same year on the y-axis. *R*_*t*+2_ is obtained from (3)–(4).
- Quadrant I: The parent-progeny relationship. Here, we have *R*_*t*+2_ on the y-axis and *A*_*t*_ on the x-axis.

In addition to Paulik diagrams, we plot the time series dynamics (in number of individuals) of the different life-history stanzas over time to study the resulting population dynamics.

### 3.3 Numerical scenarios

Given the lack of current computational approaches to replicate especially non-linear (e.g., periodic, oscillatory) SR-relationships, we focus here on three numerical scenarios:

**Scenario I**: a traditional (monotone) SR function,

**Scenario II**: a periodic (i.e., spiral) pattern, and

**Scenario III**: an oscillatory pattern.

All three scenarios are simulated individually by the DTMM presented in Section 2 and given by Eq.s (1)–(10).

Table 2 summarizes all model parameters and initial conditions used in the numerical experiments. In all scenarios, we consider the same numbers of fish at time 0. The following parameters are kept constant: the proportion *p*_*k*_ of females, probabilities *s*_*i*_ of surviving stage *i* ∈ {*e, l*, 0, 1}, and competition *β*_1_ at age 1. Furthermore, natural and fishing mortality rates for adults (with parameters *c*_*μ*_, *μ*_*u*_, *μ*_*l*_, *c*_*m*,1_, *c*_*m*,2_, *c*_*m*,3_, and *m*_*l*_) are the same for all three scenarios. Variations between Scenario I and Scenario II concern competition for larvae and juveniles of age 0 (*β*_*l*_ and *β*_0_), cannibalism (*c*_*i,k*_), and fecundity *f* (represented by parameters *f*_*u*_, *ρ*_1_, *ρ*_2_, *A*_*f*_, *c*_1_, *c*_2_, *c*_3_). In Scenario III, we use another value for the coefficient for cannibalism of eggs (*c*_*e,k*_) than in Scenario II.

**Table 2:** Parameter values and initial condition used in numerical experiments, with *k* = 2, …, 20.

| Valid for all scenarios |  |  |  |  |  |  |  |  |  |  |  |
| --- | --- | --- | --- | --- | --- | --- | --- | --- | --- | --- | --- |
| $J_{0,0}$ | $J_{1,0}$ | $N_{2,0}$ | $N_{3,0}$ | $N_{4,0}$ | $N_{5,0}$ | $N_{6,0}$ | $N_{7,0} \dots, N_{20,0}$ | | | | |
| $10^6$ | 461390 | 193687 | 21393 | 10377 | 8641 | 3341 | 1657/14 | | | | |
| $p_k$ | $s_e$ | $s_l$ | $s_0$ | $s_1$ | | | | | | | |
| 0.6 | 0.025 | 0.1 | 0.15 | 0.21 |  |  |  |  |  |  |  |
| $c_\mu$ | $\mu_u$ | $\mu_l$ | $c_{m,1}$ | $c_{m,2}$ | $c_{m,3}$ | $m_l$ | $\beta_1$ | | | | |
| 2 | 0.8 | 0.1 | 0.5 | 5.85 | 0.1 | 0.2 | 0 |  |  |  |  |
| Scenario I |  |  |  |  |  |  |  |  |  |  |  |
| | | $\beta_l$ | $\beta_0$ | $c_{l,k}$ | $c_{j,k}$ | $c_{e,k}$ | $f(k) = f_u$ | | | | |
| | | $10^{-8}$ | 0.005 | 0 | 0 | $10^{-2}$ | 25000 | | | | |
| Scenario II |  |  |  |  |  |  |  |  |  |  |  |
| $\beta_l$ | $\beta_0$ | $c_{l,k}$ | $c_{j,k}$ | $c_{e,k}$ | $\rho_1$ | $\rho_2$ | $A_f$ | $c_1$ | $c_2$ | $c_3$ | $f_u$ |
| $10^{-6}$ | $10^{-10}$ | $10^{-3}$ | $10^{-3}$ | $10^{-3}$ | 0.5 | 0 | 12 | 0.5 | 0.8 | 0.3 | 22017 |
| Scenario III |  |  |  |  |  |  |  |  |  |  |  |
| $\beta_l$ | $\beta_0$ | $c_{l,k}$ | $c_{j,k}$ | $c_{e,k}$ | $\rho_1$ | $\rho_2$ | $A_f$ | $c_1$ | $c_2$ | $c_3$ | $f_u$ |
| $10^{-6}$ | $10^{-10}$ | $10^{-3}$ | $10^{-3}$ | 0.01 | 0.5 | 0 | 12 | 0.5 | 0.8 | 0.3 | 22017 |

The three distinct sets of assumptions given in Table 2 define three Scenarios for SR-relationships and Paulik diagrams. In the following, we describe the model equations (1)–(4) illustrated in Paulik diagrams under the assumptions for Scenario I, II, and III.

#### Scenario I

For Scenario I, we assume that there is cannibalism of eggs (*c*_*e,k*_ = *c*_*e*,2_ = 10^−2^ for *k* = 2, …, 20), competition of larvae and juveniles (*β*_*l*_, *β*_0_ *>* 0), no cannibalism of larvae or juveniles, and all parameters, including fecundity, are constant with respect to *k* ≥ 2. Parameter values are given in Table 2. In this case, the DTMM equations (1)–(4) can be simplified to (11)–(13). By substituting (12) into (13) and then (11) into (12), we obtain (14). For Scenario I, the emergent SR relationship is not just any pattern, but recruitment at time *t* + 2 is a function of *A*_*t*_ for all *t* ≥ 0. The SR function is given by (14).

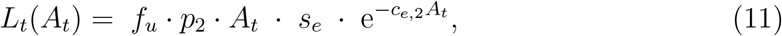

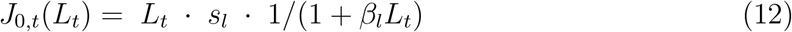

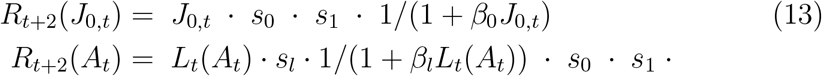

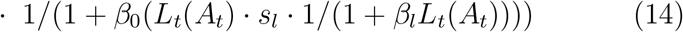

#### Scenario II

For Scenario II, we assume cannibalism of all prerecruit stages (*c*_*e,k*_, *c*_*l,k*_, *c*_*j,k*_ *>* 0 for *k* = 2, …, 20) and competition for larvae and juveniles of age 0 (*β*_*l*_, *β*_0_ *>* 0). Fecundity is a dome-shaped function of age. Under these assumptions and with parameters values as given in Table 2, the Paulik diagram illustrates the model equations (15)–(17). We observe that since fecundity is a function of age, we need the age structure of the adult population to find the number of larvae, juveniles, and ultimately, recruitment.

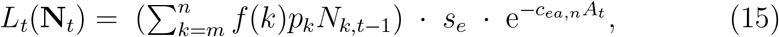

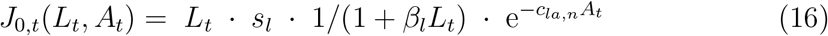

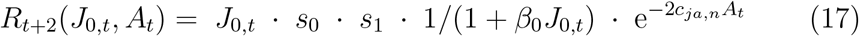

#### Scenario III

Scenario III is based on the same assumptions as Scenario II, i.e., there is cannibalism of all prerecruit stages (*c*_*e,k*_, *c*_*l,k*_, *c*_*j,k*_ *>* 0 for *k* = 2, …, 20) and competition for larvae and juveniles of age 0 (*β*_*l*_, *β*_0_ *>* 0), and fecundity is a dome-shaped function of age. The Paulik diagram for Scenario III also illustrates the model equations (15)–(17).

However, for Scenario III, we choose *c*_*e,k*_ = 0.01, for *k* = 2, …, 20, while for Scenario II, we use *c*_*e,k*_ = 10^−3^, for *k* = 2, …, 20. In Scenario III, adults have a larger effect on the survival of eggs than in Scenario II.

#### Numerical implementation of scenarios

For the numerical scenarios, we implemented (1)–(10) in MATLAB version R2020a. The difference equations are solved at 52 time steps. This allows us to see the asymptotic behavior of the dynamical system while keeping the number of runs rather low. As we are interested in the asymptotic behavior of the dynamical system, we exclude the first 10 time steps from illustrations. The solutions are illustrated using Paulik diagrams and time series plots.

## 4. Results

In this section, we present the numerical solutions of the DTMM for the three Scenarios described in Section 3.3. To study the emerging SR-relationships, we plot the numerical solutions of (1)–(10) in Paulik diagrams as presented in Section 3.2. Herein, equations (11)–(17) give a theoretical underpinning of the relationships presented in the Paulik diagram.

In the illustrations, we adopt the axes to the observed range of the solutions. The four axes in the Paulik diagram represent the intervals [*X*_*i,l*_ − 0.3(*X*_*i,u*_ − *X*_*i,l*_), *X*_*i,l*_ + 1.2(*X*_*i,u*_ − *X*_*i,l*_)], *i* ∈ S, where *X*_*a,l*_ = min_*t*=11,…,49_ *A*_*t*_, *X*_*a,u*_ = max_*t*=11,…,49_ *A*_*t*_, *X*_*l,l*_ = min_*t*=11,…,49_ *L*_*t*_, etc. A visual separation of the axes represents that 0 is not included in the axes.

### Scenario I

Figure 2 is the Paulik diagram for Scenario I. The adult-larvae relationship for the solution of the DTMM is illustrated in Quadrant IV. The underlying function given by (11) is dome-shaped, but for the range of the solution, we observe an increasing adult-larvae relationship. The larvae-juvenile relationship illustrated in Quadrant III is increasing. This relationship is described by (12) and thus an increasing function (for any domain). The juvenile-recruitment function in Quadrant II defined by (13) is increasing, too.

**Figure 2:**
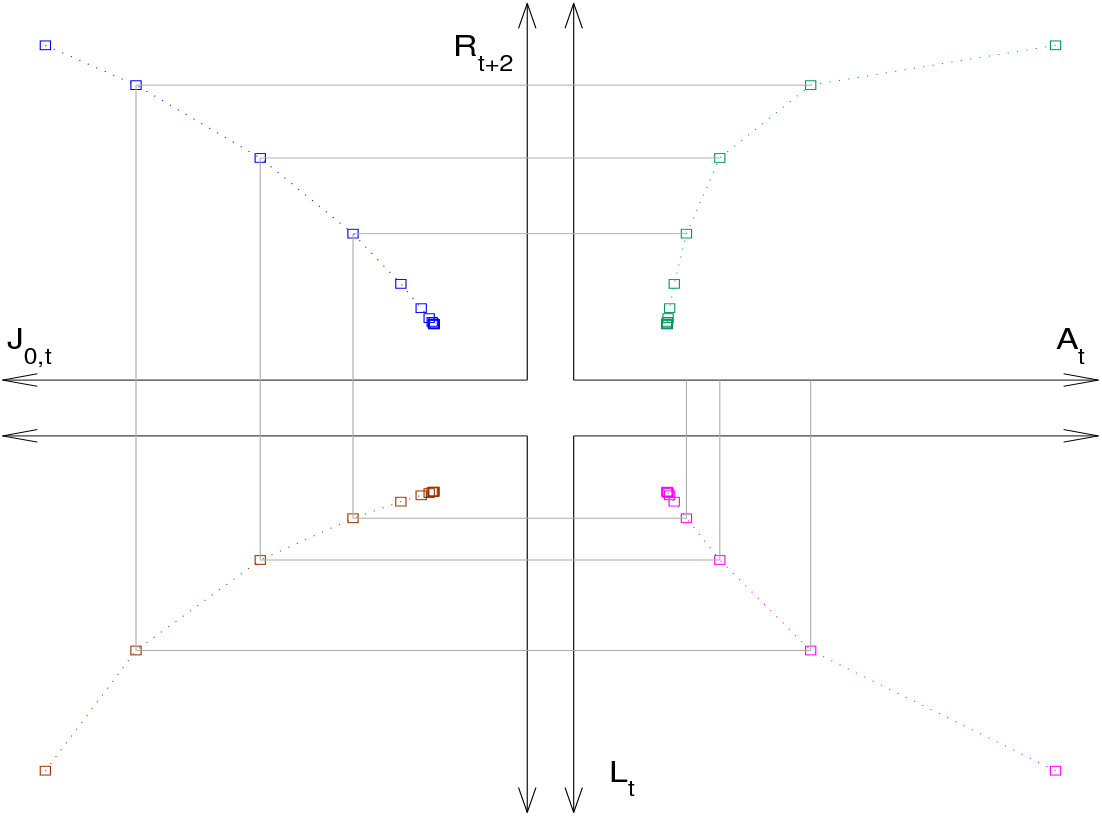
Paulik diagram of the solution of the DTMM for Scenario I. Each quadrant illustrates the number of individuals in a (sub-) stage given the number of fish in a previous stage. Dashed lines connect solutions of the dynamical system in successive years. Grey lines connect 3 instances of the number of adults and the number of larvae, juveniles, and recruits produced by them

The SR relationship shown in Quadrant I results from the relationships visualized in Quadrants II, III, and IV. For the range of the solution, the observed SR relationship is increasing. As described in 3.3, we obtain a closed-form parent-progeny function, which is given by (14). The observed SR-relationship is comparable to the typical Beverton-Holt functional form [56].

In all four quadrants, we observe that the numbers of individuals decrease over time and approach asymptotic values. This indicates that for Scenario I, the solution of the difference equation approaches an equilibrium point.

### Scenario II

The solution of the DTMM under Scenario II is illustrated in Fig.s 3– 6. The four figures illustrate the variations in the numbers of individuals over time. For Scenario II, we observe oscillatory damped behavior in the population dynamics with time.

**Figure 3:**
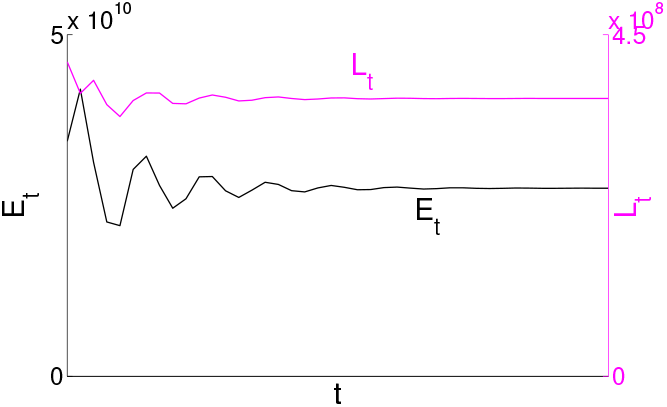
Dynamics of numbers of eggs *E*_*t*_ and larvae *L*_*t*_ for Scenario II

**Figure 4:**
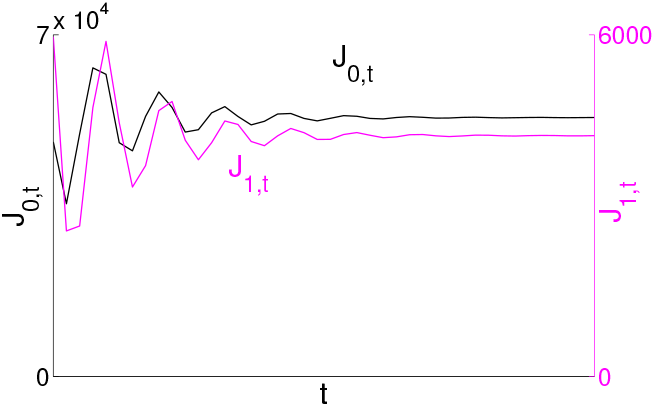
Dynamics of numbers of juveniles of ages 0 and 1 (*J*_0,*t*_ and *J*_1,*t*_) for Scenario II

**Figure 5:**
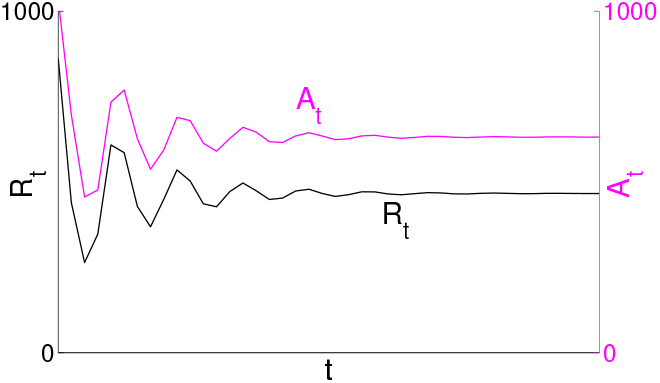
Dynamics of numbers of adults *A*_*t*_ and recruitment *R*_*t*_ for Scenario II

**Figure 6:**
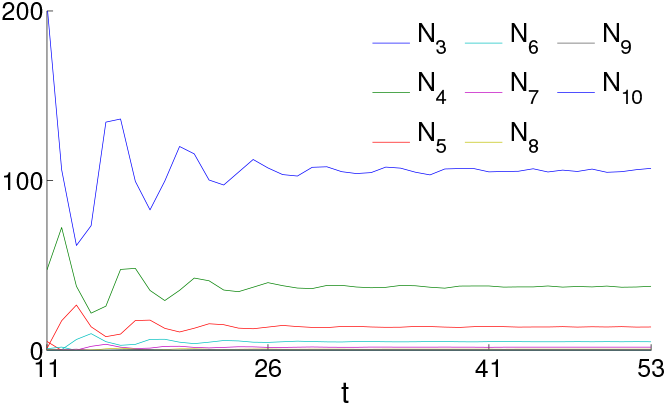
Dynamics of numbers of adults *N*_*k,t*_ of ages *k* = 3, …, 10 for Scenario II

The asymptotic behavior of the solution of the DTMM can also be observed in the Paulik diagram 7 for Scenario II. The oscillatory pattern of the SR relationship in Quadrant I results from the spiral patterns in Quadrants II, III, and IV. None of the relationships adult-larvae, larvae-juveniles, or juvenile-recruitment appear to be of closed form, and consequently, the SR relationship is not a function.

We can explain this behavior in the following way. The relationships illustrated in the Paulik diagram are given by (15)–(17). The number of larvae *L*_*t*_ defined by (15) and illustrated in Quadrant IV is in general not a function of *A*_*t*_, but of *N*_2,*t*_, …, *N*_20,*t*_. In Quadrant III, we consider the relationship between *L*_*t*_ and *J*_0,*t*_, while (16) defines *J*_0,*t*_ as a function of *L*_*t*_ and *A*_*t*_. In Quadrant II, we consider the two stages *R*_*t*+2_ and *J*_0,*t*_, but (17) defines a three-dimensional relationship between *J*_0,*t*_, *A*_*t*_, and *R*_*t*+2_. The emergent SR relationship is not a function, but the result of processes involving several stages and time steps. Overall, the observed oscillatory SR relationship under Scenario II is a result of tracking through the life cycle and is explained by the asymptotic behavior illustrated in Fig.s 3–6.

### Scenario III

In Fig.s 8–11, we illustrate the solution of the DTMM under Scenario III. Scenario III has the same assumptions as Scenario II, except for a larger value for the coefficient for cannibalism attributed to eggs. In this case, the numbers of individuals in each (sub-)stage appear to oscillate without a decrease in the amplitude with time. The solution of the DTMM oscillates but is never equal to zero.

**Figure 7:**
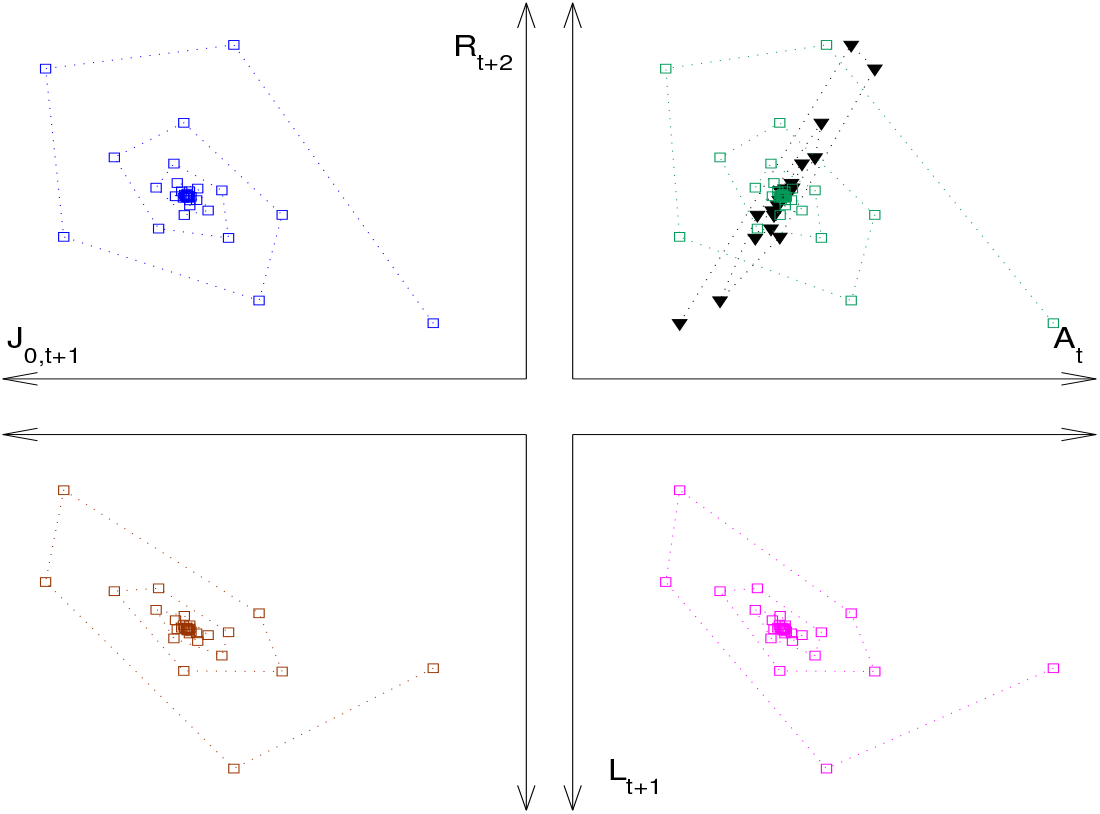
Paulik diagram of the multi-stage population dynamic system for Scenario II. Each quadrant illustrates the number of individuals in a (sub-) stage given the number of fish in a previous stage. The number of recruits *R*_*t*+2_ as a function of the number of adults *A*_*t*_ is illustrated using squares and the increase of the total number of adults *A*_*t*_ due to recruitment *R*_*t*_ with triangles

**Figure 8:**
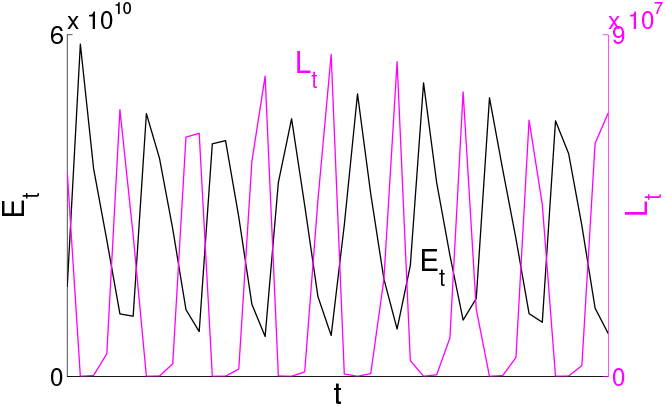
Dynamics of numbers of eggs *E*_*t*_ and larvae *L*_*t*_ for Scenario III

**Figure 9:**
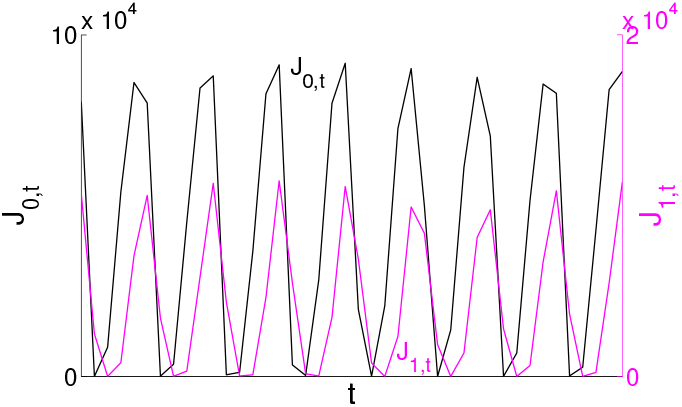
Dynamics of numbers of juveniles of ages 0 and 1 (*J*_0,*t*_ and *J*_1,*t*_) for Scenario III

**Figure 10:**
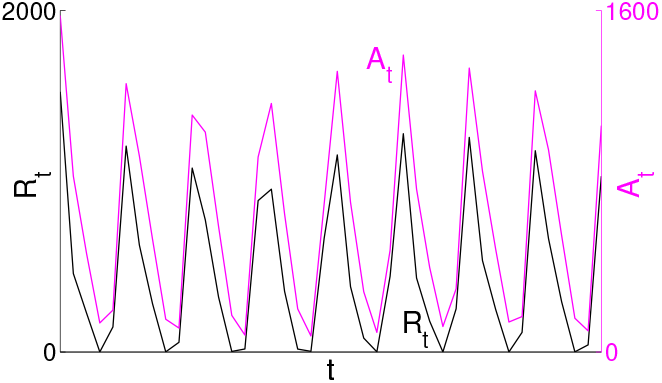
Dynamics of numbers of adults *A*_*t*_ and recruitment *R*_*t*_ for Scenario III

**Figure 11:**
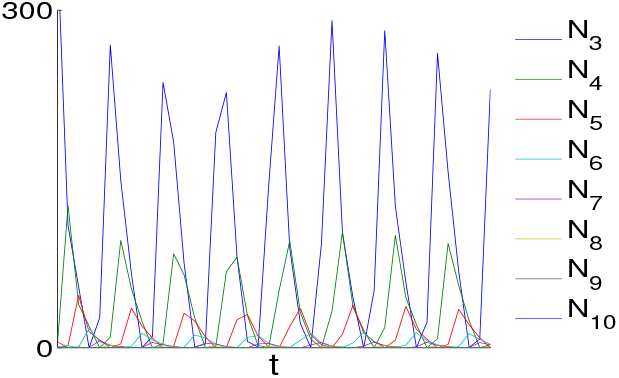
Dynamics of numbers of adults *N*_*k,t*_ of ages *k* = 3, …, 10 for Scenario III

**Figure 12:**
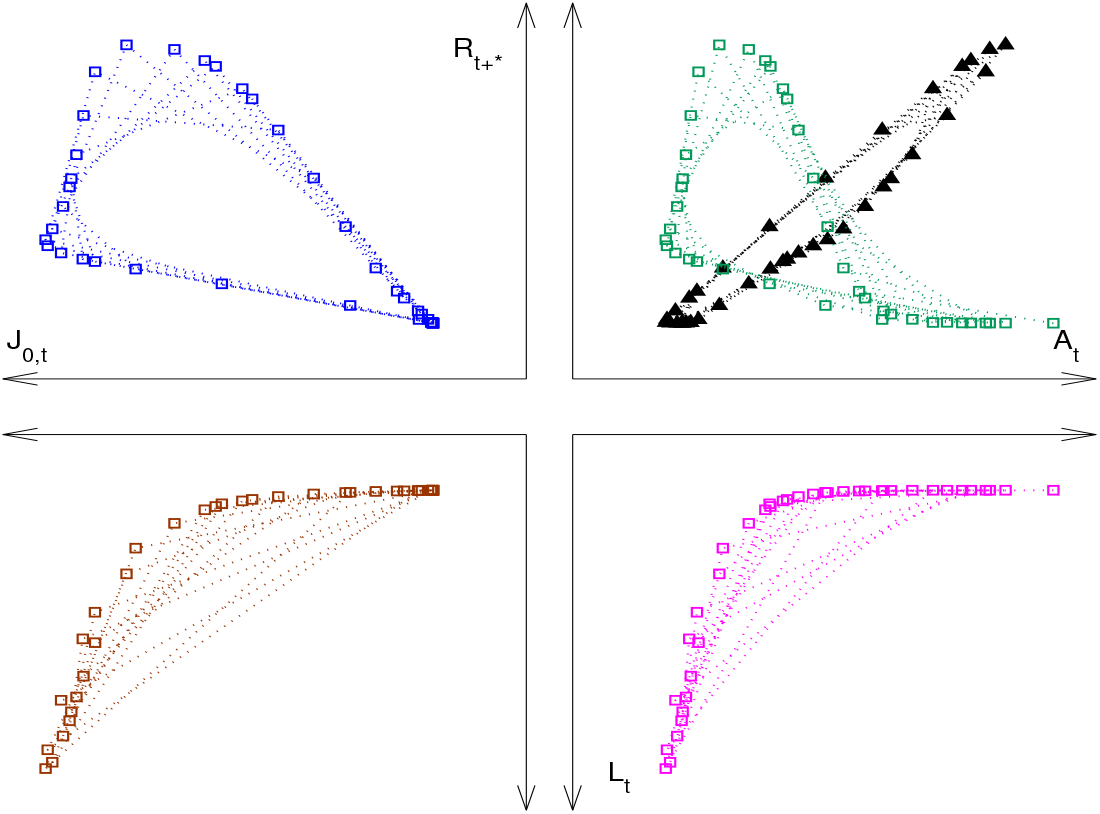
Paulik diagram of the multi-stage population dynamic system for Scenario III. Each quadrant illustrates the number of individuals in a (sub-) stage given the number of fish in a previous stage. The number of recruits *R*_*t*+2_ as a function of the number of adults *A*_*t*_ this cohort has been spawned by is illustrated using squares and the increase of the total number of adults *A*_*t*_ due to recruitment *R*_*t*_ with triangles

The Paulik diagram 12 for Scenario III illustrates the emergent SR relationship. We observe periodic patterns in Quadrants IV, III, and II, and thus, the SR relationship follows a periodic pattern. The periodic pattern cannot be captured by a closed-form SR function. As for Scenario II, this pattern can be explained by the asymptotic behavior of the DTMM.

## 5. Discussion and conclusion

This article is the first to employ a multi-stage life-cycle model to numerically simulate emergent SR relationships that are known to exist in nature.

Our model simulations reveal three specific patterns (i) a monotonic increasing trend; (ii) periodic SR patterns, and (iii) oscillatory dynamics between the stock and its recruits.

Scenarios I shows a monotone increasing pattern that is comparable to the well-known Beverton-Holt function for the SR-relationship [56]. The monotone increasing nature allows to link development of stages uniquely to one another. This characteristic observed in the Paulik diagram indicates the existence of a closed form functional SR-functional relationship (see [39]). This monotone trend in form of the the Beverton-Holt SR-function has been observed for the North Sea Herring (*Culpea harengus*) [56]. In contrast, for North Sea cod (Gadus morhua) stock the Beverton Holt funtion could only explain 10% of the variation in recruitment [32].

Scenario II shows a periodic pattern in form of spiraling inward dynamics.

The spiral inward dynamics do not allow a unique connection between the different stages, as in the case of Scenario I. In fact for a multistage model, which includes several classes of juveniles and assumes nonlinear survival functions in terms of numbers of adults, a closed form SR-function might not exist as in this case of Scenario II. Here the SR-relationship is affected by the temporal variations of the structure and state of the adult population [21].

Scenario III demonstrates how important relationships between nonsuccessive life stages are for the dynamical behaviour of the system. For the observed numbers of adults, eggs, larvae and juveniles, the relationships between adults and larvae, and larvae and juveniles, may be approximated by monotonic functions. The oscillations of the discrete time system are due to cannibalism on juveniles of both age 0 and age 1 a relationship between non-successive life stages.

In particular, empirically such periodic (oscillatory) SR patterns as in Scenario II and III, have been observed for about one half of all the cod, stocks while it was more common for slow growing individuals [31]. The authors also found that climate and temperature had a large effect on this relationship. therefore, the periodic patterns might be more profound in species that grow slower and therefore have more time to adapt to climate changes. For fast growing cod stocks the SR- patterns were rapid fluctuations [31]. The authors in [33] reported periodic (oscillatory) patterns in clock- and anti-clockwise loops for pacific stocks of Japanese sardine (*Sardinops melanostictus*), and two tuna species (*Thunnus orientalis* and *Thunnus thynnus*).

Although, we could identify SR-patterns from our scenarios also in the literature, a direct one-to-one representation is difficult especially as articles often highlight linear trends in non-linear looking SR-data (e.g., [27], or as data are logarithmic scaled on at least one axis. Several authors have however, indicated that non-linear trends in SR-patterns, similar to our results [29]. Interestingly though is that periodic (oscillatory) patterns seem often to emerge when the stock and recruitment is plotted over years. According to [57], the non-monotonic SR patterns observed in Scenarios II and III might be an explanatory factor of the oscillations commonly observed in fish population dynamics (e.g., Barents Sea Capelin (*Mallotus villosus*)) [58, 59]. Although this hypothesis needs further investigation, it displays the potential insights that can be gained by a deeper and more comprehensive understanding of existing SR patterns and their the driving factors.

The results demonstrate that by deconstructing the life-history cycle into distinct stages and developmental processes, more diverse and realistic patterns can be detected. It therefore provides a realistic representation of the dynamic relationships and feedback loops among various life-history stages and processes, surpassing the limitations of simpler analytical models.

While more comprehensive life cycle models exist in the current literature (e.g., [36, 35]), approaches that focus primarily on the SR-relationship are few and include at maximum three life stages [2, 23, 25, 26]. An exception is the DTMM approach by [39], as the first comprehensive life cycle model framework, which focuses on theoretical conditions for the existence of a closed-form SR-function. The numerical simulation approach presented here, which is based on the theory in [39], therefore fills a clear knowledge gap by presenting a comprehensive life cycle model for the SR relationship. This approach includes four life-stages eggs, larvae, juveniles and adults, and that is reconcilable with observation data. In addition, we consider an age-structured adult population and consider mortality of pre-recruits as a function of the numbers of adults. This assumption is, for example, suitable when mortality is correlated with the spawning stock biomass, as for North East Arctic Cod (*Gadus morhua*) [53].

Our DTMM is an example of the general life-cycle model described in [39] and our observations concerning the existence of a closed-form SR function under Scenarios I, II, and II can be explained by the general theory derived in [39]. For the general life-cycle model, it has been shown that a SR function may not exist, while it does exist when assuming (i) mortality of juveniles being constant with respect to the number of adults and (ii) age-independence of parameters related to egg production and survival of pre-recruits [39]. For Scenario I, we assume *c*_*j,k*_ = 0 for all *k* = 2, …, 20 and thus, mortality of juveniles is constant with respect to the number of adults. Furthermore, we assume age-independence of fecundity *f* (*k*) = *f*_*u*_ and parameters *c*_*e,k*_, *c*_*l,k*_ and *c*_*j,k*_. The existence of a SR function thus follows from the theory presented in [39]. Under Scenario II and III, both assumptions (i) and (ii) are not fulfilled and the emergent SR relationship is not a closed-form function.

Given the flexibility and adaptability of our approach, it can contribute to a more accurate assessment of a varied range of observed recruitment patterns that will lead to a better understanding of fish population dynamics. Specifically, it can (i) help identify crucial factors contributing to recruitment success by helping to recognize potential bottlenecks or critical stages with a significant impact on recruitment; (ii) offer insights into the evolution of survival strategies at different life-history stages, from predator evasion in early stages to reproductive achievement in later stages; (iii) help disentangle the complexities of fish population dynamics and life-history interactions; (iv) help investigate how environmental variability, stochastic effects, and changes in population sizes interact with different life-history stages; and (v) decipher a varied range of possible SR-patterns from specific fish species that follow a similar life cycle.

The major challenge limiting the universal usefulness of our life cycle model to the overall fisheries community and management is model calibration with currently available data. The several stages, age classes and detailed information on e.g., survival or mortality rates calls for a much greater quantity of information (data) than is currently available. We acknowledge that such information will vary with the species or stages under consideration.

In summary, we presented three SR-relationship scenarios of the more general DTMM approach by [39], a monotonically increasing trend, periodic and oscillatory dynamics. This demonstrates the applicability of our numerical approach for several discrete time models from the fisheries literature [21].Whether it is possible to recreate chaotic SR patterns, similar to chaotic population dynamics as in [34], is open for future investigation. Finally, we want to point out that our approach provides a first start in quantitatively classifying different specific SR patterns. However, as the above discussion of results has demonstrated, much more SR-patterns might exist than currently presumed. Patterns might furthermore be closely linked to e.g., subsections of specific stocks (e.g., [31]), others to climate conditions (e.g., [31]) and again others to local regions (e.g., [33]).

### Conclusion

This work bridges a critical gap in the literature by offering a holistic modeling approach that enhances our comprehension of the SR relationship, supporting improved predictions and management strategies for fish populations.

## Ethics approval

No ethical approval was needed for this study.

## CRediT authorship contribution statement

**Anna S. Frank:** Methodology, Software, Validation, Formal analysis, Investigation, Writing - original draft, Writing - review & editing, Visualization. **Ute Schaarschmidt:** Methodology, Software, Validation, Formal analysis, Investigation, Writing - original draft, Writing - review & editing, Visualization. **Richard D. M. Nash:** Conceptualization, Writing - review & editing, Investigation, Validation. **Sam Subbey:** Conceptualization, Methodology, Software, Validation, Formal analysis, Investigation, Writing - review & editing, Supervision.

## Appendix A. A general discrete time multi-stage model

Age- and stage-structured models in fisheries are commonly [22, p. 292 and Chapter 9] represented using difference equations, i.e., assuming discrete transitions between life stanzas (see e.g., Fig. 1), uniform time steps across all life stanzas, and a positive time delay between spawning and recruitment. These assumptions are consistent with the fisheries literature [22, Chapter 5]. The approach in [39] is furthermore based on the assumption that spawning and transition to the juvenile stage occur within one time step and juveniles and adults age by one in every simulation time step (if surviving). This is justified by typical stage durations of several fish populations described in [40].

The general DTMM from [39] is defined by (A1)–(A7) with initial condition (**J**_0_, **N**_0_) ∈ 𝒟 (for non-empty 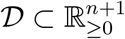 ). A nomenclature for the model is given in Table A1.

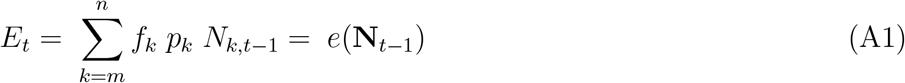

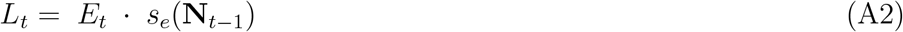

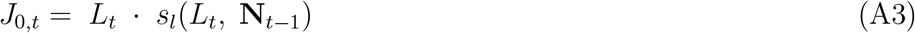

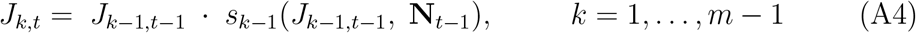

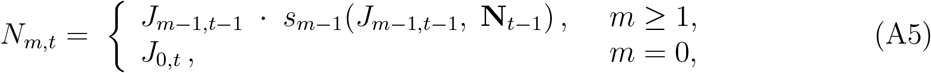

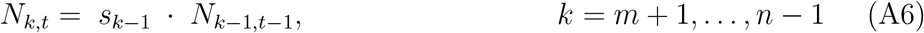

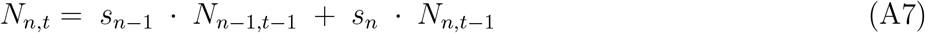

In Section 5, we refer to a result (Theorem 4) from [39] concerning the existence of a SR function. Define the following set of propositions. Conditions (C1)–(C4) are said to be true if they are true for all solutions *E*_*t*_, *L*_*t*_, **J**_*t*_, **N**_*t*_, *t* ≥ 0 of a general DTMM.

(C1) **N**_*t*_ = **N**_0_ for all *t* ∈ [0, *m* + 1],

(C2) *s*_*k*_(*J*_*k,t*_, **N**_*t*_) = *s*_*k*_(*J*_*k,t*_) for all *k* = 0, … *m* − 1 and for all *t* ≥ 0,

(C3) *m* = 0, i.e. *N*_*m,t*_ = *J*_0,*t*_ is given by (A3).

(C4)(a)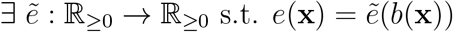, for all 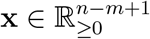,

(C4)(b)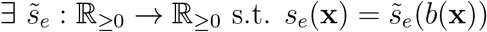, for all 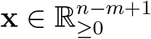,

(C4)(c) 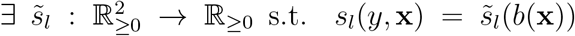 for all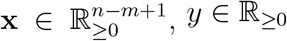,*y* ∈ **R**_≥0_.

For all general DTMMs, the logic solution of ((C4) ∧ ((C2) ∨ (C3))) ∨ (C1) is sufficient for the existence of a SR function

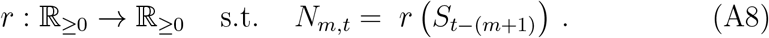

For the specific DTMM considered in Sects. 2–3, we have *S*_*t*−(*m*+1)_ = *A*_*t*−(*m*+1)_. (C2) translates to mean that (i) mortality of juveniles being constant with respect to the number of adults and (C4)(a)–(c) are implied when assuming (ii) age-independence of parameters related to egg production and survival of prerecruits. Thus, by Theorem 4 in [39], a SR function exists, if (i) and (ii) hold.

**Table A1:**
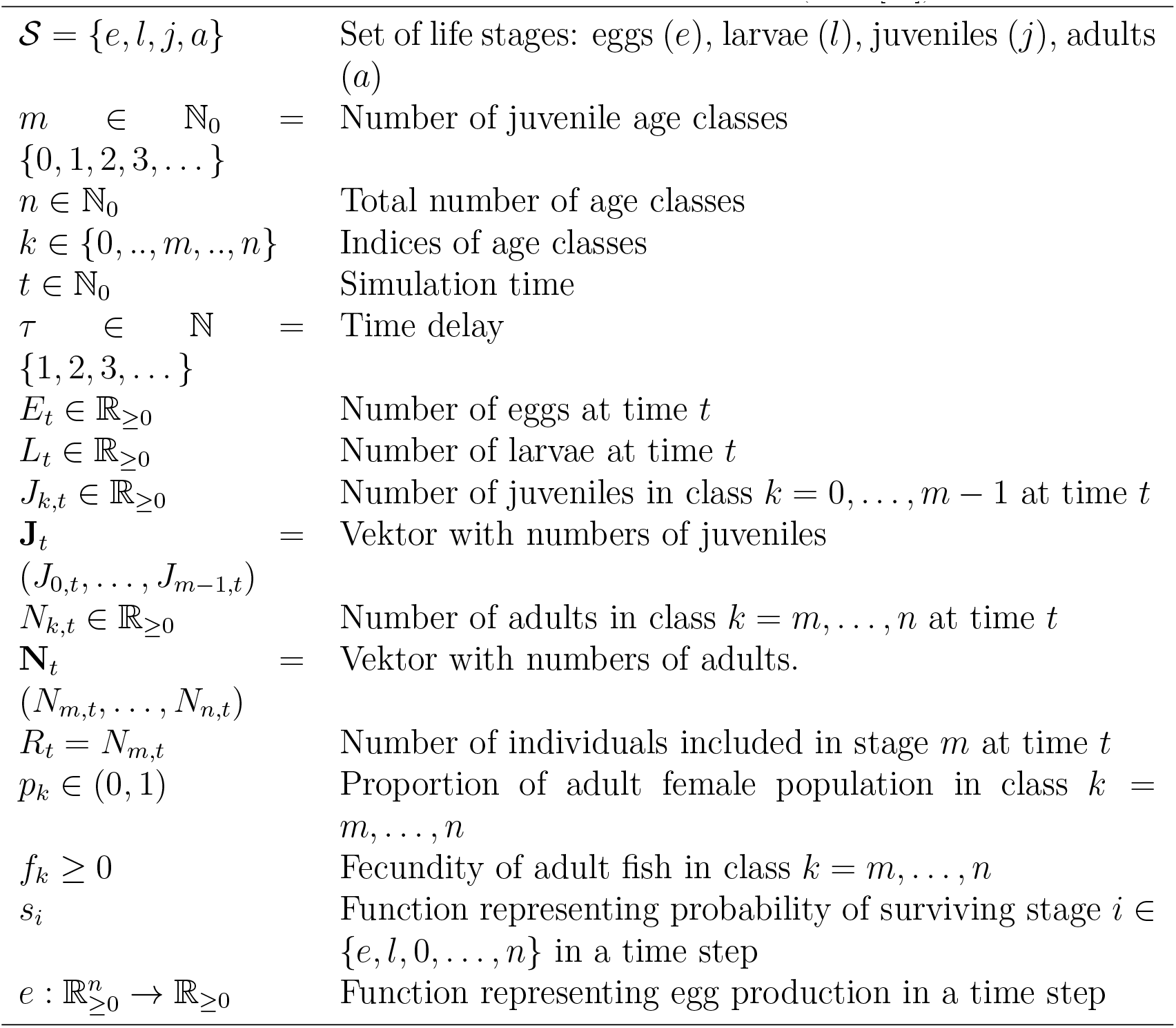
Nomenclature for the general DTMM (from [39]).

## References

[1] E. V. Camp, A. Collins, R. Ahrens, K. Lorenzen, Fish population recruitment 2: Stock recruit relationships and why they matter for stock assessment, FA234. EDIS (2021).

[2] E. N. Brooks, J. T. Thorson, K. W. Shertzer, R. D. Nash, J. K. Brodziak, K. F. Johnson, N. Klibansky, B. K. Wells, J. White, Paulik revisited: Statistical framework and estimation performance of multistage recruitment functions, Fisheries Research 217 (2019) 58–70. doi:10.1016/j.fishres.2018.06.018.

[3] E. Camp, A. B. Collins, R. N. Ahrens, K. Lorenzen, Fish population recruitment: what recruitment means and why it matters: Fa222, 3/2020, EDIS 2020 (2) (2020) 6–6.

[4] W. E. Ricker, Stock and recruitment, Journal of the Fisheries Board of Canada 11 (5) (1954) 559–623.

[5] R. J. H. Beverton, S. J. Holt, On the dynamics of exploited fish populations, Vol. 19 of Fishery Investigations. Series 2, HMSO, UK, 1957.

[6] É. E. Plagányi, M. D. Haywood, R. J. Gorton, M. C. Siple, R. A. Deng, Management implications of modelling fisheries recruitment, Fisheries Research 217 (2019) 169–184.

[7] J. Gascoigne, R. N. Lipcius, Allee effects in marine systems, Marine Ecology Progress Series 269 (2004) 49–59.

[8] K. T. Frank, D. Brickman, Allee effects and compensatory population dynamics within a stock complex, Canadian Journal of Fisheries and Aquatic Sciences 57 (3) (2000) 513–517.

[9] S. Subbey, A. S. Frank, M. Kobras, Crowding effects in an empirical predator-prey system, bioRxiv (2020) 2020–08.

[10] N. B. Goodwin, A. Grant, A. L. Perry, N. K. Dulvy, J. D. Reynolds, Life history correlates of density-dependent recruitment in marine fishes, Canadian Journal of Fisheries and Aquatic Sciences 63 (3) (2006) 494–509.

[11] G. L. Britten, M. Dowd, B. Worm, Changing recruitment capacity in global fish stocks, Proceedings of the National Academy of Sciences 113 (1) (2016) 134–139.

[12] N. Knowlton, Multiple stable states and the conservation of marine ecosystems, Progress in Oceanography 60 (2-4) (2004) 387–396.

[13] C. Möllmann, B. Müller-Karulis, G. Kornilovs, M. A. St John, Effects of climate and overfishing on zooplankton dynamics and ecosystem structure: regime shifts, trophic cascade, and feedback loops in a simple ecosystem, ICES Journal of Marine Science 65 (3) (2008) 302–310.

[14] A. Bakun, S. J. Weeks, Adverse feedback sequences in exploited marine systems: are deliberate interruptive actions warranted?, Fish and Fisheries 7 (4) (2006) 316–333.

[15] J. Gobin, N. P. Lester, M. G. Fox, E. S. Dunlop, Ecological change alters the evolutionary response to harvest in a freshwater fish, Ecological Applications 28 (8) (2018) 2175–2186.

[16] D. Nicolas, S. Rochette, M. Llope, P. Licandro, Spatio-temporal variability of the north sea cod recruitment in relation to temperature and zooplankton, PloS one 9 (2) (2014) e88447.

[17] G. Beaugrand, K. M. Brander, J. Alistair Lindley, S. Souissi, P. C. Reid, Plankton effect on cod recruitment in the north sea, Nature 426 (6967) (2003) 661–664.

[18] B. Macura, O. M. Lönnstedt, P. Byström, L. Airoldi, B. K. Eriksson, L. Rudstam, J. Støttrup, What is the impact on fish recruitment of anthropogenic physical and structural habitat change in shallow nearshore areas in temperate systems? a systematic review protocol, Environmental Evidence 5 (1) (2016) 1–8.

[19] J. X. He, D. J. Stewart, Age and size at first reproduction of fishes: predictive models based only on growth trajectories, Ecology 82 (3) (2001) 784–791.

[20] G. Ottersen, B. Bogstad, N. A. Yaragina, L. C. Stige, F. B. Vikebø, P. Dalpadado, A review of early life history dynamics of barents sea cod (gadus morhua), ICES Journal of Marine Science 71 (8) (2014) 2064–2087.

[21] U. Schaarschmidt, Multiple time–scale dynamics of stage structured populations and derivative–free optimization., Ph.D. thesis, University of Bergen (2018).

[22] T. J. Quinn, R. B. Deriso, Quantitative fish dynamics, Oxford University Press, New York, Oxford, 1999, Ch. 5.

[23] G. J. Paulik, Studies of the possible form of the stock-recruitment curve, Rapp. p.-v. réun. - Cons. int. explor. mer 164 (1973) 302–315.

[24] E. N. Brooks, J. E. Powers, Generalized compensation in stock-recruit functions: properties and implications for management, ICES J. Mar. Sci. 64 (2007) 413–424.

[25] S. Touzeau, J.-L. Gouzé, On the stock-recruitment relationships in fish population models, Environ. Model. Assess. 3 (1998) 87–93.

[26] U. Schaarschmidt, T. Steihaug, S. Subbey, A parametrized stock-recruitment relationship derived from a slow-fast population dynamic model, Mathematics and Computers in Simulation 145 (2018) 171–185, the 5th IMACS Conference on Mathematical Modelling and Computational Methods in Applied Sciences and Engineering, in honour of Professor Owe Axelsson’s 80th birthday. doi:10.1016/j.matcom.2017.10.008.

[27] T. Iles, A review of stock-recruitment relationships with reference to flatfish populations, Netherlands Journal of Sea Research 32 (3-4) (1994) 399–420.

[28] N. Caputi, A. Chandrapavan, M. Kangas, S. de Lestang, A. Hart, D. Johnston, J. Penn, Stock-recruitment-environment relationships of invertebrate resources in western australia and their link to pro-active management harvest strategies, Marine Policy 133 (2021) 104728.

[29] C. Sguotti, S. A. Otto, X. Cormon, K. M. Werner, E. Deyle, G. Sugihara, C. Möllmann, Non-linearity in stock–recruitment relationships of atlantic cod: insights from a multi-model approach, ICES Journal of Marine Science 77 (4) (2020) 1492–1502.

[30] I. G. Taylor, V. Gertseva, R. D. Methot Jr, M. N. Maunder, A stock– recruitment relationship based on pre-recruit survival, illustrated with application to spiny dogfish shark, Fisheries Research 142 (2013) 15–21.

[31] A. Rindorf, N. Cadigan, D. Howell, M. Eero, H. Gislason, Periodic fluctuations in recruitment success of atlantic cod, Canadian Journal of Fisheries and Aquatic Sciences 77 (2) (2020) 236–246.

[32] E. M. Olsen, G. Ottersen, M. Llope, K.-S. Chan, G. Beaugrand, N. C. Stenseth, Spawning stock and recruitment in North Sea cod shaped by food and climate, Proc. R. Soc. B 278 (1705) (2011) 504–510. doi:10.1098/rspb.2010.1465.

[33] K. Sakuramoto, The true mechanism that controls the stock-recruitment relationship, Open Access Library Journal 5 (2) (2018) 1–22.

[34] J. A. Wilson, J. French, P. Kleban, S. R. McKay, R. Townsend, Chaotic dynamics in a multiple species fishery: a model of community predation, Ecological modelling 58 (1-4) (1991) 303–322.

[35] E. K. Chen, N. A. Som, J. D. Deibner-Hanson, D. G. Anderson, M. J. Henderson, A life cycle model for evaluating estuary residency and recovery potential in chinook salmon, Fisheries Research 257 (2023) 106511.

[36] N. W. Kendall, J. Unrein, C. Volk, D. A. Beauchamp, K. L. Fresh, T. P. Quinn, Life cycle model reveals sensitive life stages and evaluates recovery options for a dwindling pacific salmon population, North American Journal of Fisheries Management 43 (1) (2023) 203–230.

[37] S. C. Zeug, P. S. Bergman, B. J. Cavallo, K. S. Jones, Application of a life cycle simulation model to evaluate impacts of water management and conservation actions on an endangered population of chinook salmon, Environmental Modeling & Assessment 17 (5) (2012) 455–467.

[38] F. Cordoleani, W. H. Satterthwaite, M. E. Daniels, M. R. Johnson, Using life-cycle models to identify monitoring gaps for central valley spring-run chinook salmon, San Francisco Estuary and Watershed Science 18 (4) (2020).

[39] U. Schaarschmidt, A.-S. Frank, S. Subbey, Mathematical preconditions for existence of the stock-recruitment function, Bulletin of mathematical biology (Submitted) (2023) 12pp.

[40] P. Petitgas, A. D. Rijnsdorp, M. Dickey-Collas, G. H. Engelhard, M. A. Peck, J. K. Pinnegar, K. Drinkwater, M. Huret, R. D. M. Nash, Impacts of climate change on the complex life cycles of fish, Fish Oceanogr 22 (2) (2013) 121–139. doi:10.1111/fog.12010.

[41] R. Hilborn, C. J. Walters, Quantitative fisheries stock assessment: Choice, dynamics and uncertainty, Chapman and Hall, London, 1992.

[42] J. Müller, C. Kuttler, Methods and models in mathematical biology, Springer, Berlin, 2015. doi:10.1007/978-3-642-27251-6.

[43] S. Chen, S. Watanabe, Age dependence of natural mortality coefficient in fish population dynamics, Nippon Suisan Gakk. 55 (2) (1989) 205–208.

[44] K. Siegfried, B. Sansó, A review for estimating natural mortality in fish populations, Tech. rep., SEDAR 19 Research Document 29 (2009).

[45] K. W. Shertzer, M. H. Prager, D. S. Vaughan, E. H. Williams, Fishery Models, in: S. E. Jørgensen, B. Fath (Eds.), Population Dynamics, Vol. 1 of Encyclopedia of Ecology, Oxford: Elsevier, 2008, pp. 1582–1593.

[46] G. R. Fitzhugh, K. W. Shertzer, G. T. Kellison, D. M. Wyanski, Review of size- and age-dependence in batch spawning: implications for stock assessment of fish species exhibiting indeterminate fecundity, Fish. B-NOAA 110 (4) (2012) 413–425.

[47] H. Bossel, Modeling and simulation, AK Peters/CRC Press, 2018.

[48] ICES, Report of the Working Group for the Assessment of Demersal Stocks in the North Sea and Skagerrak (WGNSSK), 30 April–7 May 2014, ICES HQ, Copenhagen, Denmark, ICES CM 2014/ACOM:13 (30 April–7 May 2014, 2014).

[49] MATLAB Release 2012a, The MathWorks, Inc., Natick, Massachusetts, United States (2012).

[50] B. J. Rothschild, Dynamics of marine fish populations, 277th Edition, Harvard University Press, London, 1986.

[51] R. D. M. Nash, Exploring the population dynamics of Irish Sea plaice, Pleuronectes platessa L., through the use of Paulik diagrams, J. Sea Res. 40 (1–2) (1998) 1–18.

[52] L. Persson, P. Byström, E. Wahlström, Cannibalism and competition in eurasian perch: population dynamics of an ontogenetic omnivore, Ecology 81 (4) (2000) 1058–1071.

[53] B. Bogstad, N. A. Yaragina, R. D. M. Nash, The early life-history dynamics of Northeast Arctic cod: levels of natural mortality and abundance during the first 3 years of life, Can. J. Fish. Aquat. Sci. 73 (2) (2016) 246–256.

[54] L. Heermann, J. Borcherding, Competition, predation, cannibalism: the development of young-of-the-year perch populations in ponds with bream or roach, Journal of Applied Ichthyology 29 (3) (2013) 549–554.

[55] D. Ricard, F. Zimmermann, M. Heino, Are negative intra-specific interactions important for recruitment dynamics? a case study of atlantic fish stocks, Marine Ecology Progress Series 547 (2016) 211–217.

[56] J. Horwood, After beverton and holt, Advances in Fisheries Science: 50 Years on from Beverton and Holt (2008) 49–62.

[57] W. E. Ricker, Stock and recruitment, J. Fish. Res. Board Can. 11 (1954) 559–623.

[58] P. A. Ahti, S. Uusi-Heikkilä, A. Kuparinen, Are there plenty of fish in the sea? how life history traits affect the eco-evolutionary consequences of population oscillations, Fisheries Research 254 (2022) 106409.

[59] A. Frank, S. Subbey, M. Kobras, H. Gjøsæter, Population dynamic regulators in an empirical predator-prey system, Journal of Theoretical Biology 527 (2021) 110814.

